# Boundary Strength Analysis: Combining colour pattern geometry and coloured patch visual properties for use in predicting behaviour and fitness

**DOI:** 10.1101/342063

**Authors:** John A. Endler, Gemma L. Cole, Alexandrea Kranz

## Abstract

Colour patterns are used by many species to make decisions that ultimately affect their Darwinian fitness. Colour patterns consist of a mosaic of patches that differ in geometry and visual properties. Although traditionally pattern geometry and colour patch visual properties are analysed separately, these components are likely to work together as a functional unit. Despite this, the combined effect of patch visual properties, patch geometry, and the effects of the patch boundaries on animal visual systems, behaviour and fitness are relatively unexplored. Here we describe Boundary Strength Analysis (BSA), a novel way to combine the geometry of the edges (boundaries among the patch classes) with the receptor noise estimate (*ΔS*) of the intensity of the edges. The method is based upon known properties of vertebrate and invertebrate retinas. The mean and SD of *ΔS (m_ΔS_, s_ΔS_)* of a colour pattern can be obtained by weighting each edge class *ΔS* by its length, separately for chromatic and achromatic *ΔS*. This assumes those colour patterns, or parts of the patterns used in signalling, with larger *m*_*ΔS*_ and *s*_*ΔS*_ are more stimulating and hence more salient to the viewers. BSA can be used to examine both colour patterns and visual backgrounds. BSA was successful in assessing the estimated conspicuousness of colour pattern variants in two species, guppies (*Poecilia reticulata*) and Gouldian finches (*Erythrura gouldiae*), both polymorphic for patch colour, luminance and geometry. The pattern difference between chromatic and achromatic edges in both species reveals the possibility that chromatic and achromatic edges could function differently. BSA can be applied to any colour pattern used in intraspecific and interspecific behaviour. Seven predictions and four questions about colour patterns are presented.

## 1. INTRODUCTION

Colour patterns are important in survival and reproduction in diverse species because they affect mating success, contests, avoiding predators, luring prey or attracting pollinators. In general, the fitness of the sender (individual with the colour pattern) is affected because the receiver (viewer of the colour pattern) can make a behavioural or physiological decision about the sender, based upon reception and perception of the sender's colour pattern (e.g. receivers will mate, fight, attack, be lured close and eaten, pollinate or disperse seeds). Colour patterns offer an effective way of investigating the complex relationship between genes, morphology, performance, fitness and evolution (Arnold 1983, 2003) because the functions of most colour patterns are relatively easy to identify (Endler 1978, 1980). However, the links between visual properties, perception, receivers' decision-making processes and fitness are not well understood.

Decisions made by the receiver depend upon both the signal design of the colour pattern (the physical structure of the signal) and its signal content (information about the signaller, reviewed in Endler 1993a). For both components, the first stage affecting fitness is the stimulation of the receiver's retina by the colour pattern; all subsequent processes leading to perception and decision making flow through this step (Lythgoe 1979). Although all components of a colour pattern may affect the viewer's decision making, their relative importance in retinal and brain stimulation is not known. In particular, we do not know how colour, luminance, patch size and patch geometry work together to affect receiver behaviour, and so cannot yet make explicit predictions about colour pattern properties or the behavioural decisions based upon them.

Relating patterns to fitness has been successful for some species with cryptic colour pattern components (Troscianko et al 2016, 2017) but there is a tendency in the literature to study only pattern or one or two colour pattern components. Previous attempts to quantify colour patterns have included mapping the pattern components (Van Belleghem et al 2018), mapping pattern component boundaries (Stevens and Cuthill 2006) and estimating the distributions of relative pattern component edge lengths (Endler 2012). Other analyses have calculated colour patch discriminability (Siddiqi et al 2004). However, all of these methods ignore whether or not the colour patches share common boundaries. Color patch boundaries are important because adjacent colour patches will influence the visual perception of a given patch as well as the contrast across the boundary.

Here we present Boundary Strength Analysis (BSA), a way to combine the effects of both patch properties and the intensity of patch edges (transitions between patches) based upon how they are processed by the visual system in the retina. BSA estimates the effects of both colour and patch edges by combining two existing methods for the first time, one for discriminability between adjacent patches (*ΔS* Vorobyev and Osorio 1998) and one for the geometric arrangement of patches (Endler 2012). Unlike all previous methods, BSA includes the estimated visual intensity of the boundaries (estimated by ΔS) and their length, rather than just recording which boundaries are present, and calculates ΔS statistics only between patches which come in contact. This is consistent with the opponent visual processes that detect colour and colour patch edges, and the fact that these processes sample small parts of the visual field (Dowling 2012, Kelber 2016). This allows us to begin to examine colour patterns less arbitrarily, by incorporating estimates of how strongly patch boundaries stimulate the retina as a proxy for conspicuousness.

BSA can be used for animal and plant colour patterns as well as visual backgrounds, and allows investigation of both within pattern and pattern-background contrast. For brevity we will describe and give examples of BSA in terms of within-pattern contrast but the resulting statistics can be calculated for visual backgrounds as well as patterns and the two compared to estimate pattern-background contrast.

### 1.2 VISUAL MODELLING OF COLOUR DISCRIMINATION

We use the receptor noise model or RN (Vorobyev and Osorio 1998; reviewed in Kelber et al 2003) to estimate detection thresholds for colour discrimination. The input to the model consists of the relative light (photon) captures for each photoreceptor class in the viewer's retina for two colour patches. The output of the RN model is *ΔS*, which is similar to a multivariate equivalent to *t* in statistics in that it compares the difference between the two sets of cone captures to the standard error of the difference. Like other signal/noise measures *ΔS* = 1 is regarded as the difference required for two colours to be noticeable, or one just noticeable difference (JND). RN predictions have been tested using behaviour of several species, and work reasonably well (e.g. Kelber et al 2003; Olsson et al 2015; Fleishman et al 2016). However, RN modelling must be used with caution for four reasons: (1) RN was designed to predict discrimination when *ΔS* is near one (near the threshold), and may be inaccurate for colours that are very different (*Δ*S > 1). This arises because the relationship between predicted difference and perceived difference is nonlinear. For example, consider three colours A, B, and C. Let the difference between A and B be *ΔS* = 2, and between A and C *ΔS* = 8; the frequent implicit assumption is that *ΔS* = 6 between B and C. Although the JND scale suggests that A and B are almost as far apart as A and C, if the perception response to *ΔS* is logarithmic then B and C may not be perceived as very different from each other and both perceived as very different from A. (2) Behaviour observations often show that some colours are discriminated as predicted by RN while others are not (unpublished observations; Cheney, pers. comm 2017; Olsson et al 2015; Fleishman et al 2016). This may arise from pre-existing colour preferences. Different RN models need to be used at higher and lower light intensities to make good predictions (Vorobyev and Osorio 1998, Olsson et al 2015). (3) Data on actual receptor noise values are scarce yet they underpin all *ΔS* calculations (Olsson et al 2017). (4) The model is limited; it is designed to capture what happens during early processing in the retina and does not include downstream processing in the brain, including decision making as well as perception. Estimates of detection and discrimination depend upon animals making decisions. Consequently, the RN could be correct in the retina, but later neural processes may mean that behaviour-tests may not match all RN predictions (for example Dyer et al 2008). Despite these limitations, what happens at the early retinal level is important because all visual processing starts there (Lythgoe 1979). The RN model must be treated simply as a starting point analogous to the Hardy-Weinberg equilibrium in population genetics. In addition to providing a foundation, RN model estimates of *ΔS* can be used to explore the visual effect of the entire colour pattern, not just differences between colour pairs.

### 1.3 ASSESSMENT OF PATCH EDGES

Previous work with colour discrimination and *ΔS* has not accounted for whether compared patches were in contact or separated by other colours. Here we explore *ΔS* explicitly for patches which come into contact because what happens at the patch edges may be important. The neurobiological justification for assessing the effects of edges (transitions between patches) is described in detail in Elder and Sachs (2004), Stevens and Cuthill (2006), Troscianko et al (2017) and Endler (2012). Briefly, the photoreceptors in both vertebrate and invertebrate visual systems are connected to neurons that calculate the differences between the photoreceptor outputs over a small visual field. Groups of photoreceptors involved in opponency are called units and can not only detect colour but also serve as edge detectors. Units consist of two adjacent groups (zones) of photoreceptors covering a small part of the visual field, and a ganglion cell calculates the difference in outputs between the two groups opponency (Dowling 2012; Dyer et al 2011; Kelber 2016; Sanes and Zipursky 2010). If the photoreceptors in the two zones are sensitive to *different* wavelengths, then the unit outputs are colour signals because colour is based upon intensity differences among different parts of the visible spectrum. Edges between patches of different colours are detected if the edge cuts across the boundary between the unit zones. If the photoreceptors in the unit are sensitive to the *same* wavelengths then the outputs result from patch edges at the zone boundary regardless of chroma if they differ in luminance. Both edge types are detected depending upon the physical size of the retinal unit relative to the image and/or how rapidly the eye scans the colour pattern (Elder and Sachs 2004; Dowling 2012; Gegenfurtner and Sharpe 1999; Kelber 2016; Sanes and Zipursky 2010). The stronger the edges (steeper gradients and greater differences between the patches, yielding larger *ΔS* between the two patches) the stronger the signal they produce in the units. The longer the edges the more units that they will stimulate. Consequently, both the geometry and reflectance spectra of patches in colour patterns affect edge intensity and conspicuousness. Both chromatic and achromatic opponent units operate over small parts of the visual field, suggesting that local colour pattern properties may be more important than global properties.

The effects of edges also depend upon the visual acuity (resolution angle) of the viewer as well as the distance between the viewer and the colour pattern. Acuity effects may eliminate or modify visual contrast, particularly if the visual fields of the opponent units are larger than the patches. Although opponent units are known to cover a small part of the visual field, their actual sizes are unknown in most species. Moreover there may be higher-order units in the brain which will not be accounted for by the retinal estimations. For these reasons, calculations of edge effects must be done with good data on acuity and viewing distance, and results treated as a first approximation, even if the unit field sizes are known.

### 2 METHODS

Let *C* be the number of colour and luminance classes in a given colour pattern. The challenge of this, and any other colour pattern analysis, is identifying the *C* classes and making identification repeatable. This is a classic image analysis problem known as image segmentation, and is particularly problematic where there are colour or luminance gradients. One could identify the classes by (human) eye, but for almost all diurnal nonprimate animals their vision is sufficiently different from humans that human-based classifications may range from unreliable to misleading, particularly if there are UV reflecting patches present. Another method is to move a portable reflectance spectrometer sensor over the animal's body to determine how patch reflectance spectra vary. If any of the spectra vary more than is visible to the human eye then samples must be taken from both the invisible and visible patches and labelled accordingly. A third method which is less likely to miss patches invisible to humans is to scan the entire body evenly in a grid with a spectrometer and use various clustering methods to classify the colour/luminance patches by spectral clusters. This can be refined by doing clustering of calibrated photographic pixels (Van Belleghem et al 2018), spectra or cone stimulations and clustering based upon *ΔS* (van den Berg et al, in preparation). A final stage is ensuring that all patches in the segmented image are visible with the viewer’s visual acuity and viewing distance. In what follows, we will assume that the patch classification into *C* classes has been completed along with a matching list of cone captures estimated from patch spectra (Endler & Mielke 2015) or from calibrated photographs (Troscianko and Stevens 2015).

All cone capture estimates should be made under the normal viewing conditions in the wild. This includes the distances between signals and receivers as well as light intensity because visual acuity declines with declining light and the combination of the visual acuity of the viewer and the viewing distance affects the smallest patch which can be resolved. If two patches are not resolved at the ordinary distance and light intensity, then the two patches should be combined into a single patch and the patch spectrum should be an average of the two spectra, weighted at each wavelength by the relative areas of the two indistinguishable patches. The geometry of patches should be relevant to the viewer’s vision and visual conditions during viewing.

### 2.1 RELATIVE FREQUENCY OF EACH PATCH EDGE CLASS

The first stage of analysis of a colour pattern is to estimate the lengths or relative frequencies of the *C* edges between adjacent colour/luminance patch classes. A *C* × *C* matrix should be made to organise the list of all possible edge or colour/luminance transition classes (example in supplemental table S1). For *C* classes there are at most *E = C*(*C*-1)/2 different edge or transition classes (Endler 2012). Note that in any one colour pattern it is likely that not all patch classes will contact all other classes, especially for larger *C*. Consequently, the number of observed kinds of different transitions (edges) among patches, *n*, will be less than the maximum possible number of edge classes, E. A simple example is found in the North American coral snakes *Micrurus fulvius* and *M. euryxanthus*, where there are colloquial phrases to distinguish them from the Batesian mimetic king snakes *(Lampropeltis)* such as "red on yellow, beware the fellow, red on black, it's all right Jack". There are three possible transitions in these snakes: red-yellow, yellow-black and red-black, but red-black is a missing transition in these coral snakes while and red-yellow is missing from the mimics (this is not true for other coral snake species). Once the edge classes are determined, they need to be mapped onto the outline of the animal. An example using a male guppy *(Poecilia reticulata*) is shown in Fig. 1A-C.

**FIGURE 1.**
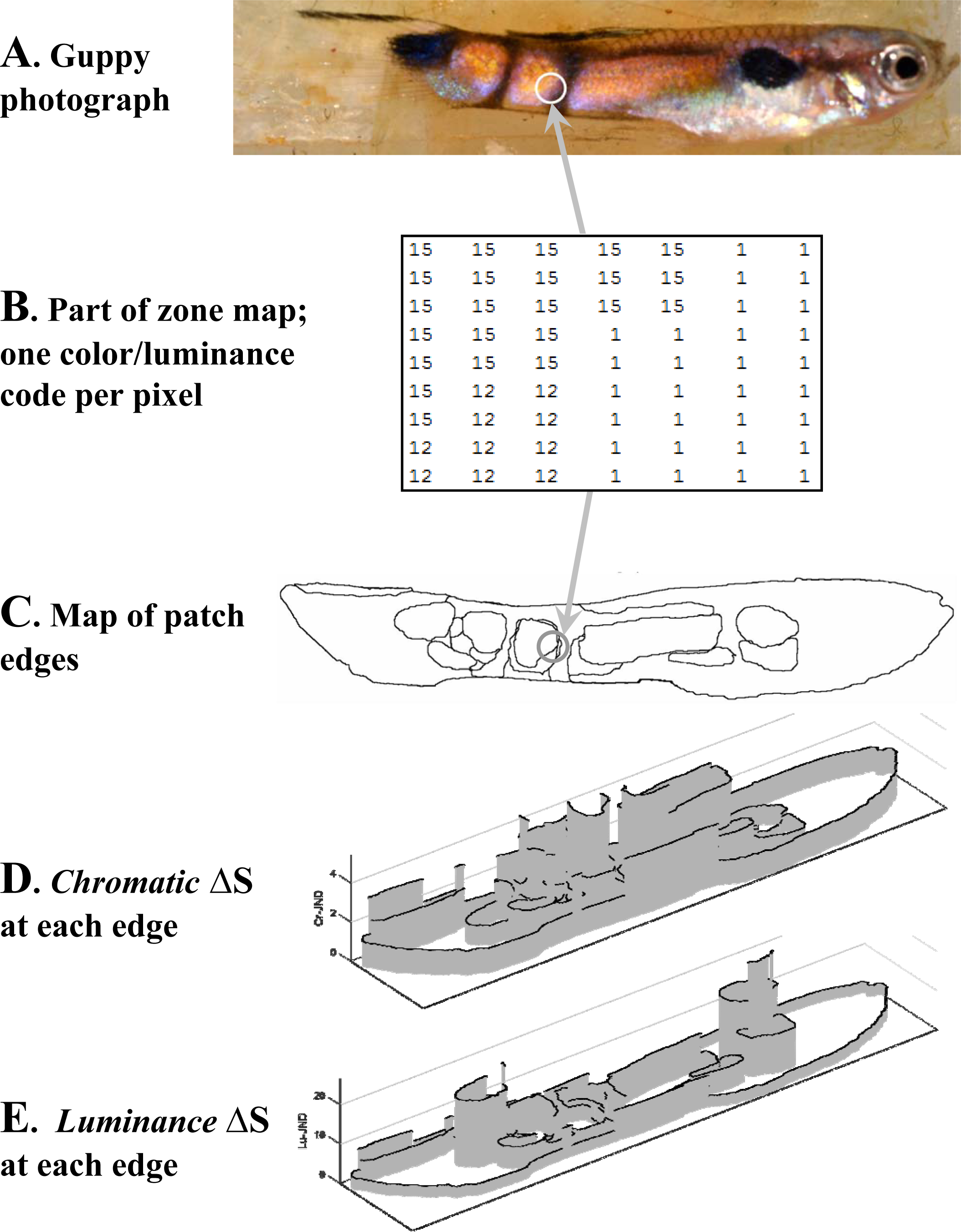
Example analysis of a male guppy colour pattern. (A), photograph of a guppy (scale not shown). (B), part of the resulting zone map indicated by the circle in panels (A) and (C). Each pixel has a code indicating which colour/luminance class overlaps that pixel (see Endler 2012 for details). (C) edge map; this can either be derived directly from the photograph (A) or from the zone map (B). (D), Diagram in which the x,y (horizontal) coordinates correspond to the edge map in (C) and the vertical axis corresponds to the chromatic *ΔS* between adjacent patches under specific ambient light conditions. (E) as in (D) but for luminance *ΔS*. Note the lack of topographic correspondence between the chromatic and luminance diagrams. For brevity we will refer to (D) and (E) as “Fort Diagrams” because they resemble old fashioned fortresses).

The relative frequency or length of each transition class can be obtained from one of two methods. Measure the length for each edge directly from the edge map (Fig. 1C) or extract edges from the zone map of the patch pattern. A zone map is simply a digital mosaic diagram of the same size as the original image where each pixel contains a label for the colour/luminance class in which it is found (Fig. 1B); this is also known as a label matrix. The zone map also allows additional parameters to be extracted (Endler 2012). Because pixels are in a square array, diagonal distances as well as horizontal or vertical distances will have to be used for slanted edges, but this should produce minor errors if the pixel spacing is small enough. Accumulating the colour/luminance class transitions over all adjacent pixels in the zone map yields a transition or adjacency matrix, where rows and columns correspond to the colour classes (as in table S1). The transition matrix diagonal entries are proportional to each colour's relative area. The off-diagonals yield the relative frequency of each transition class or edge (Endler 2012). This matrix is symmetric with separate estimates of a particular transition class in both the upper and lower off-diagonals (table S1). For further analysis, add the equivalent upper and lower off-diagonals together in order to obtain frequencies of each patch edge type (table S2); these numbers are equivalent to lengths of edges extracted directly from the image (Fig. 1C), and, like lengths, can be divided by their grand total to yield relative edge lengths. The result of either method is a *C* × *C* lower offdiagonal transition or edge matrix, **T_E_** (table S2), where the lower off-diagonal numbers are the lengths or frequencies of the edge class defined by the intersection of the corresponding row and column. For example, if a particular cell (row *i* and column *j)* has the value *f*_*ij*_, then *f*_*ij*_ is the frequency of the transition between colours *i* and *j* in both directions. Potential transitions between colours which are not observed because the appropriate patches do not come in contact will be represented by *f*_*ij*_ = 0. A given *f*_*ij*_ in **T_E_** estimates how commonly two colour/luminance classes share a common edge or the size of each patch type boundary.

### 2.2 MAGNITUDE HENCE SALIENCE OF PATCH BOUNDARIES

The second and novel stage of analysis is an estimate of how conspicuousness the edge is likely to be to a given viewer under given environmental conditions. The receptor noise *ΔS* estimate for any pair of colours is an estimate of edge conspicuousness or strength because colour and/or luminance differences are easier to detect for larger *ΔS*. We can obtain photon captures for each patch using the irradiance spectrum illuminating the pattern in nature, the reflectance spectrum of the patch in the direction of the viewer, the transmission spectrum of the air or water between the pattern and viewer in nature, the transmission spectrum of the eye optics, and the absorption spectra of the visual pigments in each photoreceptor class (Lythgoe 1979; Endler & Mielke 2005; Kelber et al 2003). We obtain the *ΔS* for all possible pairs of patches in the colour pattern (as did Siddiqi et al 2004) based upon the photoreceptor captures, the relative abundance of each photoreceptor, and an assumption about the level of receptor noise (the Weber fraction, Kelber et al 2003). Methods for obtaining *ΔS* are well established, including in the R package *pavo* (Maia et al. 2013). The *ΔS* for each kind of colour class comparison is then placed in the appropriate row and column in a second matrix with *C* rows and *C* columns (same format as table S2). It is only necessary to fill in the lower off -diagonal because the upper off diagonal should be identical, and the diagonals will be zero (no difference in a comparison of the same colour). This yields a *C* x *C* transition or *ΔS* matrix **T_S_** with data in the lower off-diagonal, where each entry *s*_*ij*_ is the *ΔS* for patch colour/luminance classes indicated by row *i* and column *j*. Two different **T_S_** should be calculated by: (1) using all the photoreceptors used in colour vision (e.g. cones in vertebrates) to obtain chromatic *ΔS* and (2) using the specific photoreceptor(s) used in luminance to get luminance or achromatic *ΔS*. Consequently the result will be two *ΔS* transition matrices, **T_SC_** from the chromatic *ΔS* calculations and **T_SL_** from the luminance or achromatic *ΔS* calculations. Ensure that the rows and columns of **T_SC_** and **T_SL_** correspond exactly in both length (*C*) and row order to the rows and columns of **T_E_.**

The matrix **T_E_** should contain the relative frequencies of each kind of transition and the matrices **T_SC_** and **T_SL_** Should contain the RN estimate of how differently (*ΔS*) the two adjacent colours in the corresponding **T_E_** entry stimulate the retina with respect to chromaticity or luminance, respectively. They should have the same form as table S2. The lower off-diagonal values of these three matrices should be converted to vectors (one-dimensional lists) of length *E = C*(*C*-1)/2, and placed together in a *E* × 3 data matrix for convenience in further calculations (see table S3). This data matrix has the edge length, the chromatic *ΔS*, and the luminance *ΔS* for the transition (edge) class *k* in row *k*; call these *f*_*k*_, *sc*_*k*_ and *sl*_*k*_ where *k* = 1 ̤ *n* patch classes. Table S3 shows an example where *k = a,b̤.f* and n=9.

The data matrix provides a correspondence between edge lengths and their estimated visual magnitudes or salience. This, along with an annotated map of the patch boundaries (Fig. 1C), allows plotting the geometry of estimated patch boundary strengths for both chromatic and luminance *ΔS*. In these diagrams the x and y axes are as in Fig. 1C and the z-axis is proportional to *ΔS*. Fig. 1D,E show 3D plots of chromatic and luminance edge *ΔS* for the guppy shown in Fig. 1A. We will call these diagrams “fort diagrams” because they resemble forts and “fort” means strong in French and Latin, so also refers to boundary strength. Note the very different geometric patterns of chromaticity and luminance boundaries in Fig. 1 D and E; the guppy shows high edge contrasts in different places for chromaticity and luminance. More specifically, luminance contrast is dominated by the black patch edges almost independent of the patch class they contact. Note the very high luminance *⊿S* (height) where a black patch contacts the very highly reflective silver patch towards the front of the guppy in (compare Fig. 1 A and E).

### 2.3 COMBINING PATCH PROPERTIES AND EDGES

If edges contribute significantly to the conspicuousness of the entire colour pattern, then we may be able to capture at least part of what makes a colour pattern conspicuous by obtaining an aggregate measure of the edge magnitudes. We suggest the mean, standard deviation and CV of the edges’ *ΔS*, weighted by their corresponding lengths or frequencies. These are calculated from either the *sc*_*k*_ (chromatic *ΔS)* or *sl*_*k*_ (luminance *ΔS)* as *s*_*k*_ from

**T_SC_** or **T_SL_**, and using the *f*_*k*_ (from **T_E_**) as weights in the formulae:

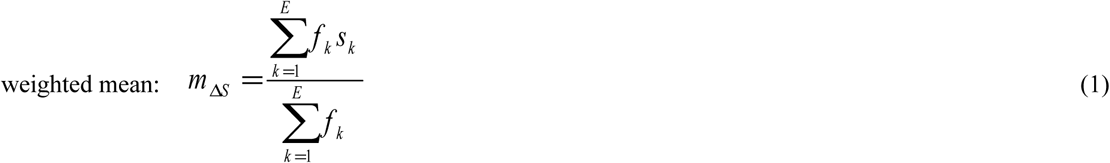

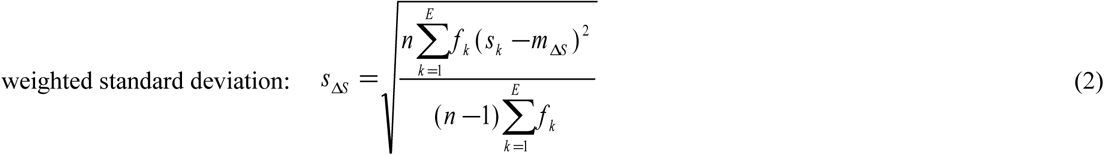

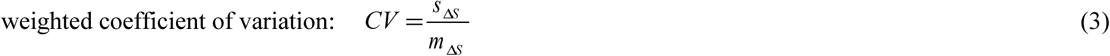

where E is the number of all possible different kinds of edges and *n* is the number of observed transitions or those with non-zero *f*_*k*_ (Filliben et al 1996); *n≤E.* The supplemental appendix provides a MATLAB function to calculate the weighted mean and standard deviation; the equivalent functions in R are wt.mean and wt.sd within the R package *SDMTools* (Van der Wal et al 2014). Formulae 1 - 3 are the same formulae used to calculate the mean, SD and CV of chroma and luminance for overall within-contrast measurements, substituting chroma or luminance for *s*_*k*_ and mean chroma or luminance for *m*_*ΔS*_; but circular statistics have to be used for hue angles (Endler & Mielke 2005).

The weighted mean *m*_*ΔS*_ is an estimate of the average conspicuousness of the whole pattern but weighting longer edges more than shorter ones. Similarly, the weighted standard deviation *s*_*ΔS*_ measures how variable the edge magnitudes are over the entire pattern weighted by their lengths. The coefficient of variation CV is the standard deviation relative to the mean. If it is known that the viewer attends only to part of the pattern then *m*_*ΔS*_ and *s*_*ΔS*_ should be calculated over the relevant part of the colour pattern. The assumption here is that a longer edge will stimulate more opponency units in the retina, and when the pattern is moving, a longer edge will sweep out more of the retinal area than a smaller edge. It is not known or obvious whether the mean, standard deviation, or even the CV would be a better predictor of salience. For example, a larger *m*_*ΔS*_ might be more stimulating, but it is unknown whether this should be accompanied by a smaller *s*_*ΔS*_ for consistently high stimulation over the entire pattern, or a larger *s*_*ΔS*_ and hence less predictable edge magnitude to prevent sensory adaptation. Using CV instead of the standard deviation might be important if a given degree of variation is not more important for small versus larger means. These conjectures can only be answered by extensive behavioural studies with different *m*_*ΔS*_ and *s*_*ΔS*_, measured under the appropriate conditions and appropriate parts of the body.

Boundary Strength Analysis (BSA) can be applied to an animal colour pattern in order to estimate within-pattern visual contrast. They can also be applied to visual backgrounds to estimate within-background contrast, and if so estimates of signal-background contrast can be made by comparing parameters of animal and background. For simplicity the results will concentrate on within-signal contrast.

## 3.0 EXAMPLES AND THEIR IMPLICATIONS

To illustrate and explore the biological significance of BSA, we chose two species that are polymorphic in their patch colour, luminance and geometry, male guppies *(Poecilia reticulata)* and Gouldian finches *(Erythrura gouldiae*) because they have very different signal and signalling geometry. This allows us to showcase the power of the method in colour pattern research and the important effects of local patterns and viewing angle between the sender and receiver.

### 3.1 GUPPY EXAMPLES AND IMPLICATIONS

Male guppies are extremely polymorphic in patch geometry and properties (Endler 1978, 1980). Fig. 2 shows Fort diagrams of six male guppies in the same format as Fig. 1C, D, ordered by decreasing chromatic *m*_*ΔS*_ and calculated in open/cloudy light conditions (Endler 1993b). The numbers are *m*_*ΔS*_ and CV from equations (1) and (3). These six randomly selected guppies yield five observations: (1) Each guppy has edges with unique geometry. This goes with the considerable polymorphism of male guppy colour patterns (photos in Endler 1978). (2) There is little geometric correspondence between the strength and positions of chromatic and achromatic (luminance) edges; the peaks in chromaticity do not correspond with peaks in luminance, and both depend upon which pair of patches form the edge. The spatial correlation between chromatic and luminance *ΔS* is always negative within a guppy although not always significantly so (Fig. 3A,B). (3) The negative correlation between the two *ΔS* is not present when we consider all possible patch combinations (Fig. 3C); patch contacts and hence boundary strengths are clearly non-random. (4) Guppies differ in how variable their *ΔS* heights are, indicating variation in which patches form common edges.

**FIGURE 2.**
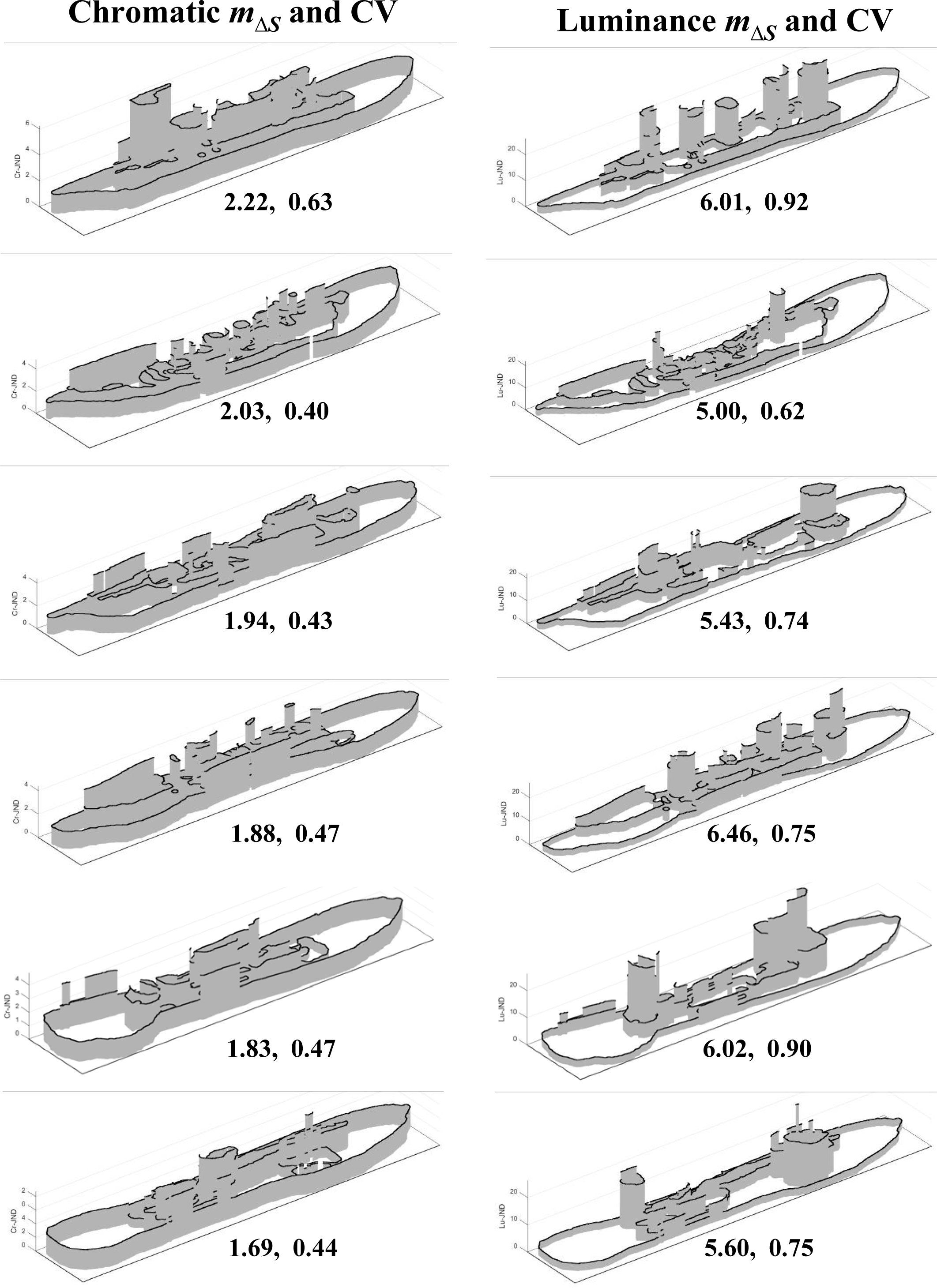
Examples of Fort Diagrams for 6 different guppy colour patterns, arranged in order of decreasing chromatic *m*_*ΔS*_. Rows correspond to the same individual guppy and columns refer to the guppy's chromatic or luminance Fort diagram, respectively. Numbers under the diagrams for each row are chromatic *m*_*ΔS*_ and CV (left column) and luminance *m*_*ΔS*_ and CV (right column) for the same guppy. Note the lack of topographic correspondence between the chromatic and luminance diagrams, and the variation among individuals.

Maximum chroma and luminance should be negatively correlated because the only way to increase chroma is to remove parts of a spectrum. Removing part of the spectral radiance reduces luminance. At the same time it increases the differences in stimulation among different photoreceptor classes, increasing chroma (Endler and Théry 1996; Endler and Mielke 2015). However, *m*_*ΔS*_ and *s*_*ΔS*_ depend upon geometry as well as patch properties and consequently predictions based upon patch properties alone may be invalid. For example, chromatic and luminance *m*_*ΔS*_ might even be positively correlated if sexual selection jointly increases both luminance and chromatic *m*_*ΔS*_, which would make males more conspicuous. We tested for a possible chromatic-luminance relationship by analysing 200 male guppies. The two *m*_*ΔS*_ are positively correlated (Fig. 3D). This is not what one would expect from random patch geometry, where every patch class has an equal probability of contacting the others (see also Fig. 3A,C). It suggests that particular colours are adjacent and adjacency has evolved to set particular levels of overall conspicuousness, as estimated by *m*_*ΔS*_. Random associations yield different *m*_*ΔS*_. The relationship for *s*_*ΔS*_ is also positive (Fig. 3E), but the 200 points are widely scattered and appear in 3 clumps. This suggests partially discontinuous variation among fish boundary *ΔS*, and could result from polymorphic colour pattern genes that control particular sets of spots (review in Endler 1978). The correlation and clumping for CV (Fig. 3F) is lower than for *m*_*ΔS*_ and *s*_*ΔS*_. Patterns of variation in boundary strength could predict fitness in any species because they affect pattern conspicuousness and hence colour pattern function and fitness.

**FIGURE 3.**
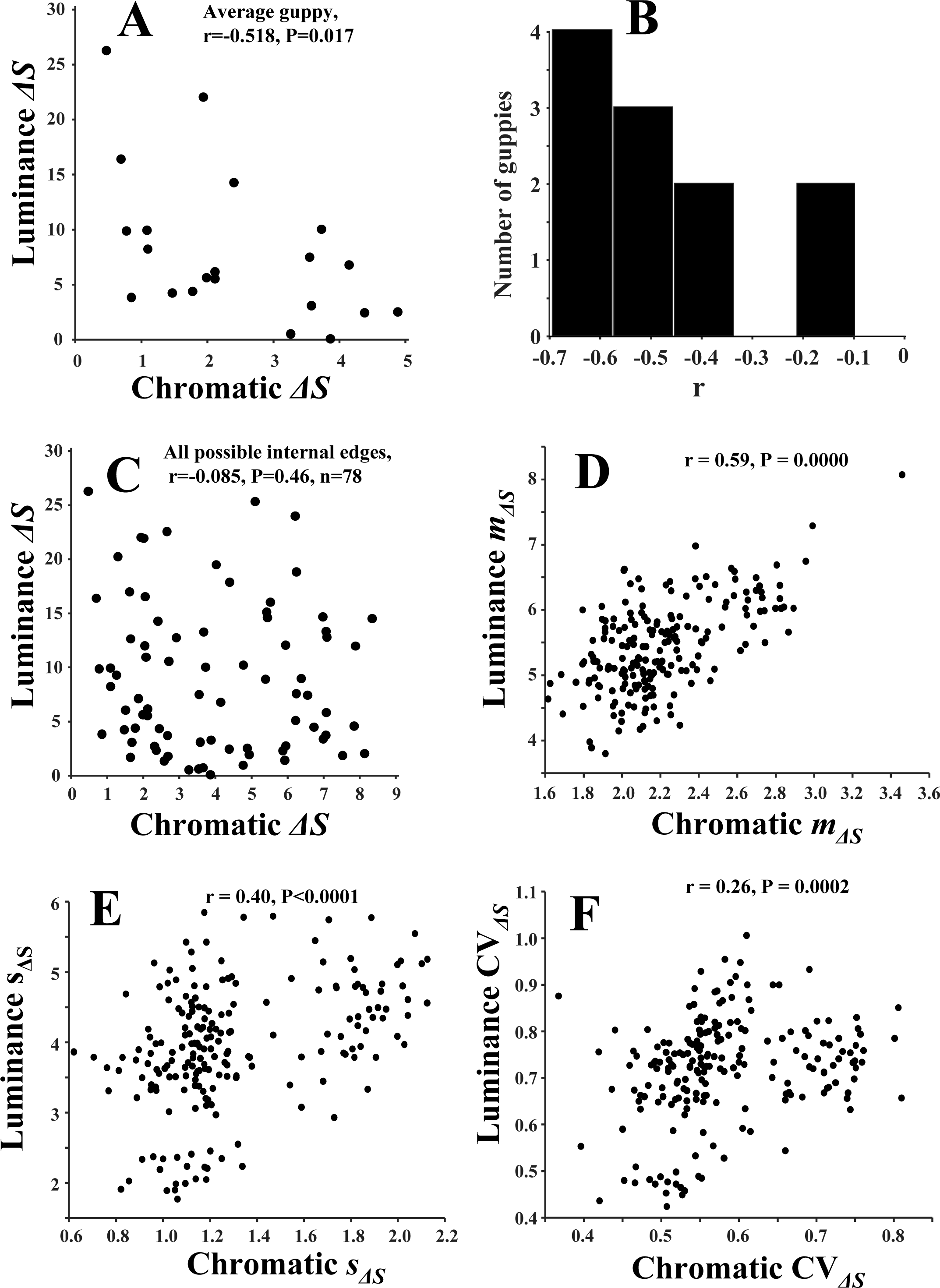
Relationships between chromatic and luminance in guppies. A, Significant negative correlation between chromatic and luminance *ΔS* within a guppy having an average correlation value. B, Distribution of the correlations among 11 guppies; all are negative but two are not significantly negative. C, Lack of correlation between all possible chromatic and luminance edges; note the larger rage and higher joint values compared to A. D, The relationship between chromatic and luminance *m*_*ΔS*_ of 200 guppies. E, Relationship for *s*_*ΔS*_. F, relationship for CV_*ΔS*_

Fig. 4 shows chromatic and luminance *m*_*ΔS*_ and *s*_*ΔS*_ distributions for the 200 guppies analysed. The means are moderately symmetrically and unimodally distributed but the standard deviations are multimodal, as in Figs. 3E, F. Note that *m*_*ΔS*_ > 1.5 indicates that, on average, the boundaries are detectable by females, but some may not be *(m*_*ΔS*_ = 1 is one JND, the threshold for distinguishing patches). Patches with similar colours or luminances which would lead to smaller *ΔS* and *m*_*ΔS*_ tend not to be adjacent. In general, we hypothesize that having adjacent patches with larger *ΔS* would be advantageous in conspicuous signalling, but disadvantageous for crypsis. If most boundaries are not detectable and a few were, this might be a previously unrecognised form of disruptive colouration.

**FIGURE 4.**
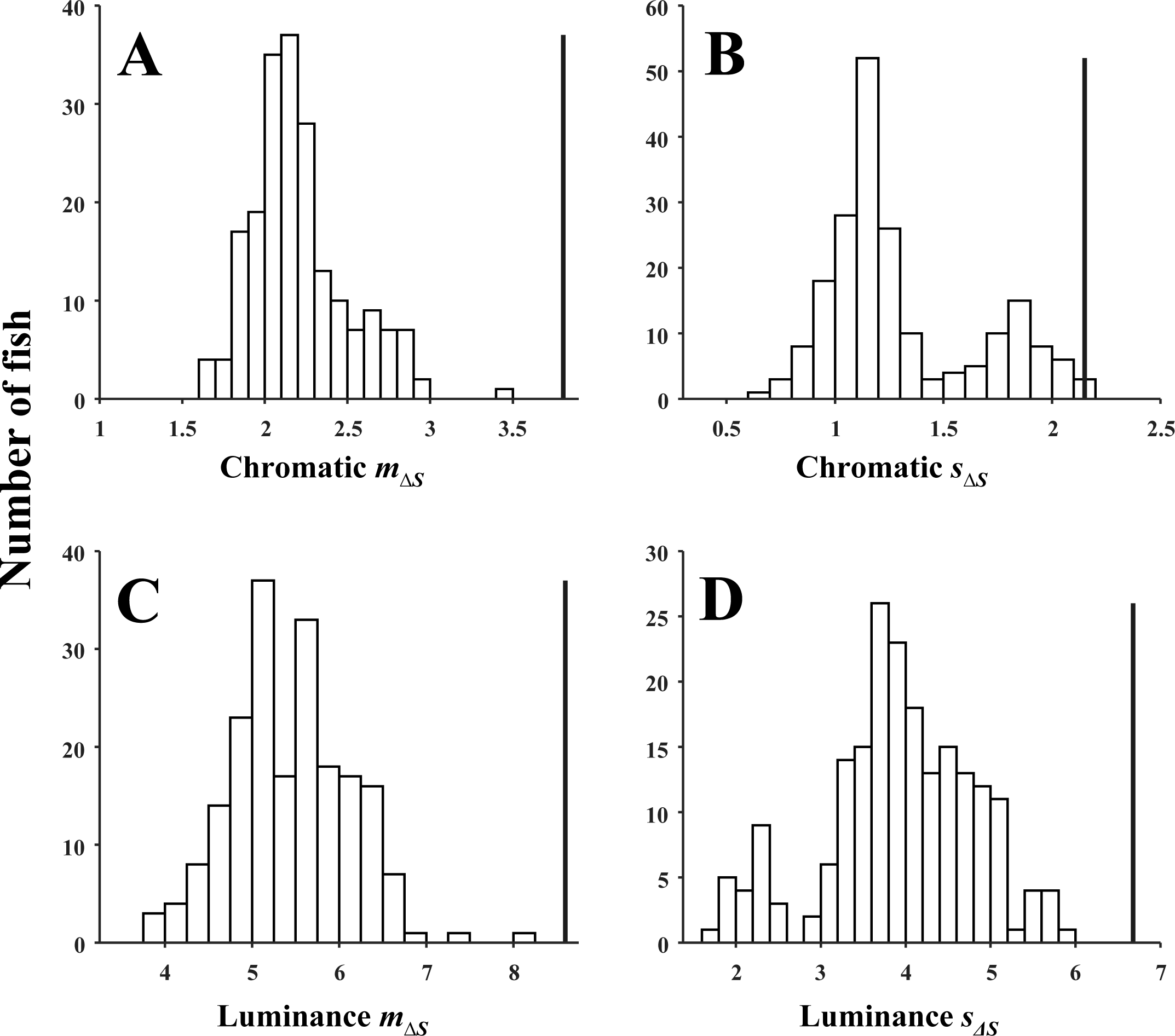
The distributions of chromatic and luminance edge statistics *m*_*ΔS*_ and *s*_*ΔS*_ of the 200 guppies in Figs. and 4. (A), chromatic *m*_*ΔS*_, (B), chromatic *s*_*ΔS*_, (C), luminance *m*_*ΔS*_, (D) luminance *s*_*ΔS*_. All guppies have *m*_*ΔS*_ >1 indicating that adjacent patches are always discriminable to guppies under the environmental conditions. The thick vertical lines show the same statistics if the colour patches were distributed at random over each guppy's body; every patch class had an equal probability of contacting the others. Almost all guppies show smaller values than expected from random patch locations.

The thick black line in Fig. 4 is the estimate for randomly arranged patch classes, as opposed to their observed geometry. This was calculated by letting every patch class contact every other patch class as in Fig. 3C. For *m*_*ΔS*_ it is larger than actually found in any fish, and for *s*_*ΔS*_ it is larger than all fish except for chromatic *s*_*ΔS*_ where it is larger than 98% of the fish. This suggests that the observed colour patterns are less conspicuous than they would be if the patches were arranged at random. One would at first think that this is contrary to that expected because we assume that females should mate with males with larger *m*_*ΔS*_ because they are more conspicuous than those with smaller *m*_*ΔS*_. However, visually hunting predators are always present in natural guppy populations, resulting in variation in the trade-off between sexual selection and predation (Endler 1978, 1980). We speculate that guppies have been selected over millions of generations for optimal edge strengths balancing sexual selection and predation. We predict that samples taken from high predation populations would have distributions of *m*_*ΔS*_ and *s*_*ΔS*_ that extensively overlap *ΔS*=1, indicating less conspicuous coloration representing the local balance between sexual selection and predation. This may apply to any species where there is a shifting balance between sexual selection and predation.

### 3.2 GOULDIAN FINCH EXAMPLES AND IMPLICATIONS

Gouldian finches provide examples of additional insights that can be gained from Boundary Strength Analysis. There are three polymorphs differing in head colour: black, yellow (golden) or red. Both males and females are coloured with females having less chromatic colours and a mauve rather than a purple chest. Unlike guppies, which have a relatively flat surface that is displayed towards females, Gouldian finches have a 3D colour pattern in which the relative proportion of patches and edges changes with viewing angle. Consequently we present Fort diagrams from Gouldian finches seen at two viewing angles: a¾ view and a side view (Fig. 5A, B). The analysis of the ¾ view is shown in Figs. 5 and 6 and the side view in Fig. 7. More details are shown in the online appendix.

**FIGURE 5.**
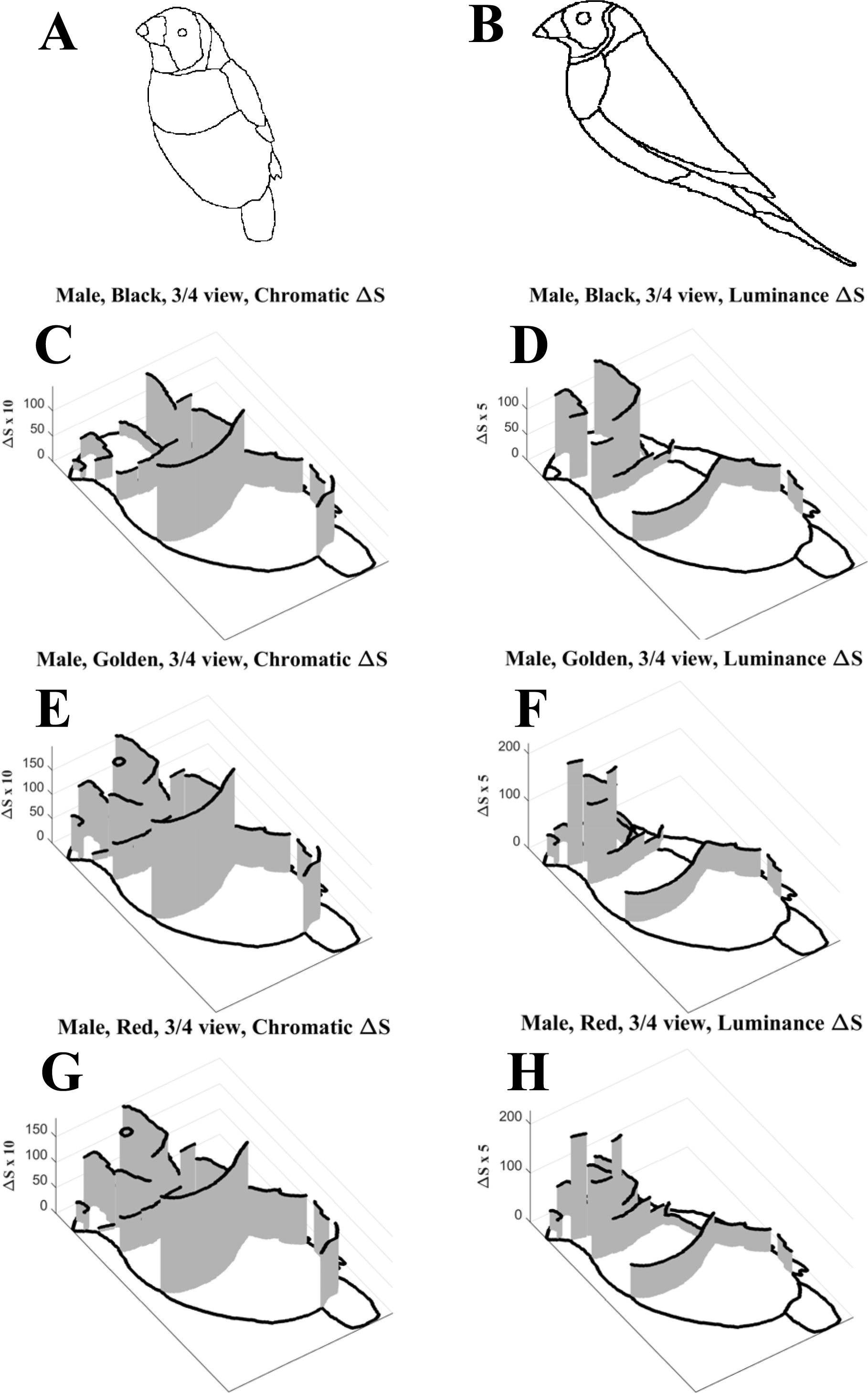
Gouldian finches. A, edge map traced from a 3/4 view photograph. B, edge map traced from a side view photograph. C-H, fort diagrams of the three male morphs (rows) showing the difference in pattern for chromatic and luminance *ΔS* (columns) in the 3/4 view.

**FIGURE 6.**
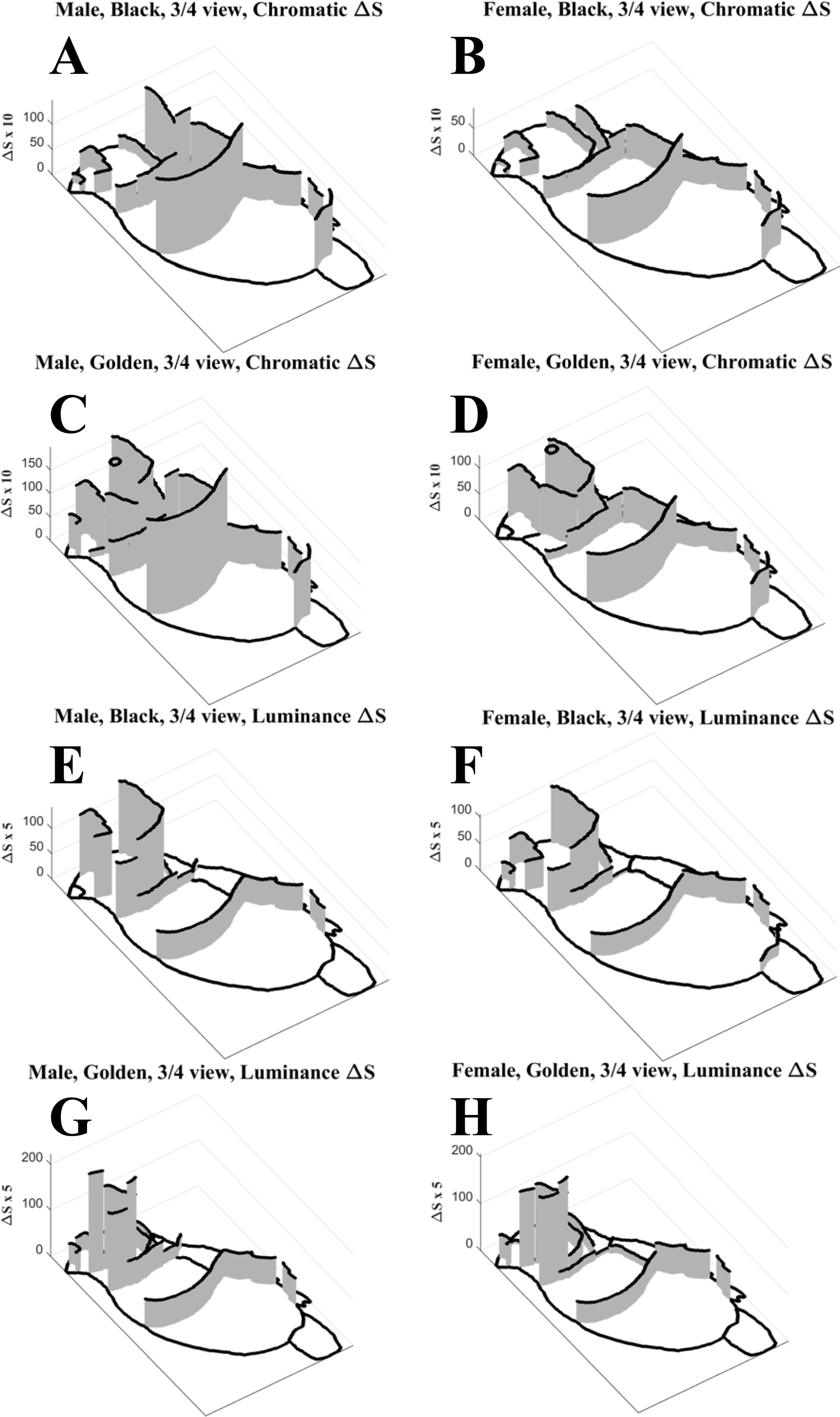
Fort diagrams showing sexual dimorphism in the black and golden-headed morphs with respect to both chromatic and luminance *ΔS* in the 3/4 view. The red-headed morph does not differ very much from the golden-headed morph (see online appendix for all fort diagrams).

**FIGURE 7.**
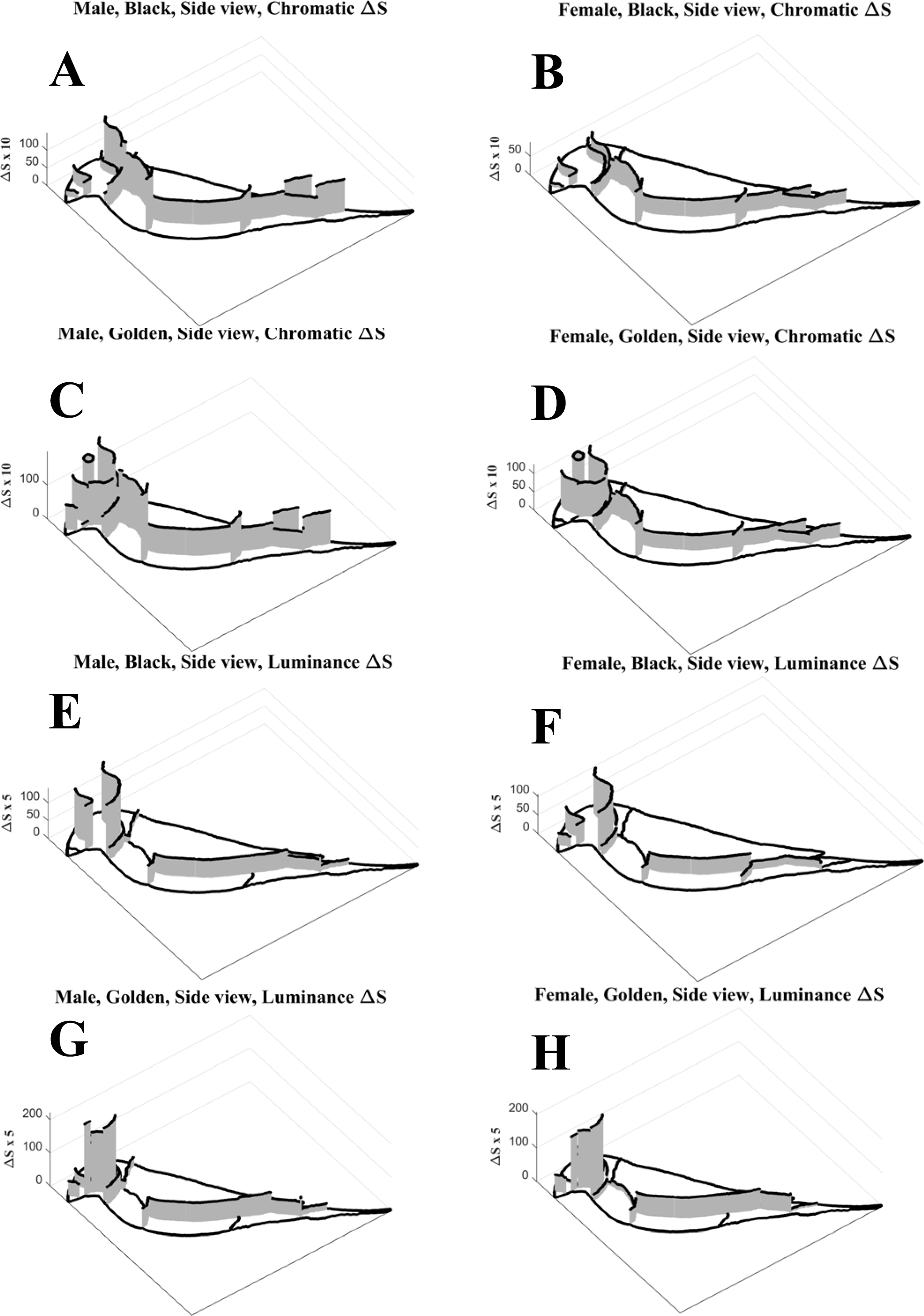
Fort diagrams of side views of the black and golden-headed morphs. See online appendix for all fort diagrams.

Like guppies, there is a divergence between chromatic and luminance *ΔS* (Fig 5C-H) and the spatial correlation between them is negative (except in the golden female morph). With fewer points than in the guppy data, none of the correlations are significant. Nevertheless, each correlation is smaller than the correlation between all possible pairs of colours for that morph and gender (see online appendix) suggesting that the negative correlation has some function in both species.

Given that the chromatic and achromatic patterns are different and almost complementary we suggest that the chromatic and achromatic components of colour patterns could be used for different functions, such as sexual selection, species recognition, or defense. Chromaticity and luminance are processed independently, and there is variation in their relative importance in stimulus choice and discrimination, among many species including crabs, psyllids, honeybees, bumblebees, flies, hawkmoths, birds and humans (Baldwin and Johnsen 2012; Farnier et al 2014; Dyer et al 2008; Giurfa et al 1997; Kelber 2005, 2016; Kiel et al 2013; Osorio and Vorobyev 2005; White and Kemp 2016; White et al 2017; Zhou et al 2012). This suggests that chromatic and achromatic channels could have different functions in any taxa. There are also distance effects, probably due to the fact that in many animals, visual acuity is greater for achromatic than chromatic stimuli. For example, bees use chromatic cues when they subtend larger angles on their retina and achromatic cues when the visual angles are smaller (Giurfa et al 1997). This means that achromatic cues may be more useful at greater distances than chromatic cues, especially at lower light levels when acuity decreases, and colour vision stops working at still lower irradiances. Moreover, chromatic and luminance components are roughly independent in natural scenes (Hansen and Gegenfurtner 2009) suggesting that crypsis may be possible independently of signalling. The functional differences between chromatic and achromatic edges are worth further investigation.

Gouldian finches also illustrate that: (1) The viewing angle significantly affects the perceived relative area of each patch, significantly affecting *m*_*ΔS*_ and *S*_*Δs*_; the ¾ view having higher *m*_*ΔS*_ and often higher *s*_*ΔS*_ than the side view (Table 1). This highlights the importance of recording the viewing angle during visual signalling. (2) Sexual dimorphism within each morph is associated with reduced edge intensities, *m*_*ΔS*_ and *s*_*ΔS*_, in females of all morphs for both chromatic and achromatic *ΔS* (Fig. 6, Table 1), with less reduction in achromatic *ΔS* (Table 1). This illustrates the utility of BSA in estimating sexual dimorphism. (3) Within males or females, the three morphs differ in chromatic *m⊿S* with the golden and red morphs similar but different from the black morph (Table 1). They differ less in achromatic *m*_*ΔS*_, and there is surprisingly little variation in *s*_*ΔS*_ among morphs; perhaps this is the sign of a species-specific signal. (4) There is a clear difference in pattern between the head and the rest of the body, with the head values larger than the body. The difference in location-specific edge intensities is stronger in the side view. This reiterates the importance of calculations using the same view angle as used by the viewers, but it also shows a weakness of using *m*_*ΔS*_ and *s*_*ΔS*_ calculated over the entire body. It may be reasonable in guppies or other species that present the entire side of a relatively flat surface to the viewer, but it will be inaccurate if the viewer attends more to some parts of the body than the others. The stronger edges in the Gouldian finch heads may be associated with, and even selected by, conspecifics paying more attention to the heads than the rest of the body. The rest of the body may be used in species recognition and, or, reduction of predator risk. Consequently, *m*_*ΔS*_ and *s*_*ΔS*_ should be calculated on the parts of the colour pattern used in social interactions for signal design assessment whereas they should be calculated separately on the parts of the body seen by predators (using predator vision parameters). These two functions may be spatially separated. Clearly we need to know about the geometry of signalling as much as the geometry of the signals for accurate use of BSA.

**Table 1.** Gouldian finch mean (*m*_*ΔS*_) and SD (*s*_*Δs*_) of patch edge chromatic (Cr) and luminance (Lm) *ΔS*, weighted by edge lengths

| **Cr $m_{\Delta S}$** | **Cr $s_{\Delta S}$** | **Lm $m_{\Delta S}$** | **Lm $s_{\Delta S}$** | **Morph-gender-view** |
| --- | --- | --- | --- | --- |
| 7.56 | 4.97 | 11.07 | 10.43 | Black, Male, 3/4 view |
| 5.71 | 4.25 | 7.84 | 9.29 | Black, Male, Side view |
| 4.49 | 2.64 | 8.55 | 6.53 | Black, Female, 3/4 view |
| 3.19 | 2.21 | 5.78 | 6.50 | Black, Female, Side view |
| 12.30 | 5.46 | 11.84 | 11.08 | Golden, Male, 3/4 view |
| 8.58 | 5.55 | 9.75 | 10.91 | Golden, Male, Side view |
| 6.70 | 3.43 | 9.90 | 10.41 | Golden, Female, 3/4 view |
| 4.77 | 3.57 | 8.33 | 9.91 | Golden, Female, Side view |
| 11.44 | 4.94 | 12.95 | 9.56 | Red, Male, 3/4 view |
| 7.96 | 4.95 | 9.68 | 10.10 | Red, Male, Side view |
| 5.75 | 2.98 | 11.80 | 9.30 | Red, Female, 3/4 view |
| 4.40 | 3.25 | 9.76 | 9.24 | Red, Female, Side view |

## 4.0 GENERAL PREDICTIONS

Because BSA can be used to analyse any animal or plant colour pattern, it is useful to make some general predictions, based upon the assumption that edges are important in colour pattern detection and perception (Gegenfurtner and Sharpe 1999; Dowling 2012; Stevens and Cuthill 2016), and that stronger edges (larger ⊿S and greater length) are more effective.

1. If *m*_*ΔS*_ is important in intraspecific signalling then it should predict behaviours such as mate choice or any other visually-based choice behaviour. The relative importance of chromatic and luminance *m*_*ΔS*_ is unknown, and this may vary among higher taxonomic groups. Consequently, we predict that the relationship between *m*_*ΔS*_, pattern conspicuousness, decision-making, and fitness will be context, habitat and species specific. Restriction of *m*_*ΔS*_ to calculations just over the part of the colour pattern tracked by viewers should be limited to species with well-studied signalling geometry, or will have to wait for more advances in eye-tracking methodologies

2. If *s*_*ΔS*_ is important in colour pattern conspicuousness then it should predict visually-based choices. However, it is not clear whether larger or smaller *s*_*ΔS*_ increases the overall conspicuousness. Small *s*_*ΔS*_ (or CV) could give a consistently higher stimulation to the retina. However, larger *S*_*⊿s*_ might be more effective if spatially similar *ΔS* (low *s*_*ΔS*_) leads to sensory adaptation and hence inefficient reception.

3. For colour patterns, or components used in signalling, edges should have *m*_*ΔS*_ > 1 with respect to chromatic and luminance *ΔS;* edges with *ΔS*≤1 are unlikely to be detected. Patterns with small *m*_*ΔS*_ have fewer detectable edges, leading to inefficient visual signalling. For crypsis, having mostly undetectable edges *(m*_*ΔS*_ ≤ 1) is an advantage. However, if the background has many *ΔS*>1 and the animal has many *ΔS*≤1the animal's shape will be conspicuous. If both have many *ΔS*>1 then the pattern may be cryptic (Endler 1978) or disruptively coloured (Endler 2006).

4. For colour patterns or pattern parts used in signalling, the distribution of both *m*_*ΔS*_ and s_ΔS_ should be different from those of the visual background with respect to either chromatic or luminance *ΔS* or both. The animal-background colour pattern component distributions should be similar for cryptic species, or parts of the colour patters that are seen more often by predators than conspecifics.

5. The animal-background match or mismatch of both *m*_*ΔS*_ and s_ΔS_ should differ in different parts of the animal's body for species that are usually seen by predators from one viewing angle (e.g. above or behind) and by conspecifics from another viewing angle (e.g. frontal; e.g. Salticid spiders); parts viewed by predators should be more cryptic than parts viewed by conspecifics. Colour pattern functions could not only differ in regions of the body viewed from different angles, but may also differ when viewed from different distances because this may cause some adjacent patches to blend (Endler 1978).

6. For prey species living in areas over a range of predation intensities, the fraction of edges with *ΔS*≤1 should be relatively higher in areas with higher predation because *ΔS*≤1 leads to poorer perception of separate patches, but the opposite is needed for disruptive colouration. The absolute fraction of edges with *ΔS*≤1 should depend upon the background patch pattern. For example, in visual backgrounds with highly contrasting patches (most *ΔS* ≫1, large *m*_*ΔS*_) the *m*_*ΔS*_ and the distributions of *ΔS* in the animal and backgrounds should be more similar in areas of higher predation intensity than areas of lower predation. For prey species that use only parts of the pattern for signalling, the signalling components should be smaller, with shorter edges and lower *ΔS* in areas of greater predation risk.

7. For species attending more to chromaticity than luminance in intraspecific signalling the chromatic *m*_*ΔS*_ and most or all chromatic *ΔS* should be larger than 1 with the opposite for luminance. This ensures that the pattern is maximally conspicuous to the receiver's visual system. A similar pattern should appear for luminance *m*_*ΔS*_ and *ΔS* in species using luminance more than chromaticity.

## 5.0 GENERAL QUESTIONS

There is so little known about the implications of estimates of patch boundary strengths that predictions are limited, but there are several questions which are worth further investigation until we can make explicit predictions.

1. Which is more important in intraspecific signalling, *m*_*ΔS*_ or *s*_*ΔS*_*?* If both are important, does their relative importance change with the complexity of the visual background or the mixture of different intraspecific and interspecific viewers?

2. *m*_*ΔS*_ and *s*_*ΔS*_ estimate the effects of patch boundaries on the overall colour pattern conspicuousness. It is also possible that within-pattern variation in hue, chroma and luminance of patches also affect overall conspicuousness, regardless of whether or not they come into contact (Endler & Mielke 2005). What is the relative importance of overall variation in hue, chroma, luminance, and edge properties? Which measures successfully predict mate choice and survival under specific visual and ecological conditions?

3. Do different aspects of salience allow for “private channels”, allowing mitigation of the tradeoff between being conspicuous to potential mates and inconspicuous to predators? This might be most likely if, for example, predators used different visual processing, different components of the colour patterns, or different viewing distances than the prey use for intraspecific signalling.

4. How do patch and patch edge properties communicate signal content? Do they constrain content enough to make predictions about the kind and amount of information to be transmitted to conspecifics?

In sum, within the limitations outlined in sections 1.2 and 1.3, Boundary Strength Analysis will enable these questions to be addressed in any species that use vision to make decisions based upon reception and perception of a sender's colour pattern.

## ACKNOWLEDGEMENTS

We thank three reviewers for excellent and useful comments on the manuscript, Adrian Dyer for useful comments about the receptor noise model and Adelaide Sibeaux for comments on the manuscript and being willing to try it out as a way to predict guppy mating success (in progress). We thank the Australian Research Council for two discovery grants which supported this research (DP110101421 and DP150102817).

## AUTHOR'S CONTRIBUTIONS

JAE devised the method, tested it, and wrote the first draft of the paper. GLC and XK prepared the guppy photographs, extracted the colour patch geometry from photographs, and helped revise the paper.

## DATA AVAILABILITY

A MATLAB script for calculating weighted means and standard deviations is found in the online supplemental material.

## Supplemental information

## 1. Example of transition matrix terminology and calculations

### The Edge Matrix, **T_E_**

Consider a color pattern in which there are *C* = 4 colors, *c*_i_, where *i* =1,2,3,4. The edge matrix T_E_ can be generated either from a zone map (Endler 2012) or directly from a map of the edges.

If **T_E_** was generated from a zone map, the raw **T_E_** data will appear as in Table S1.

**Table S1.**
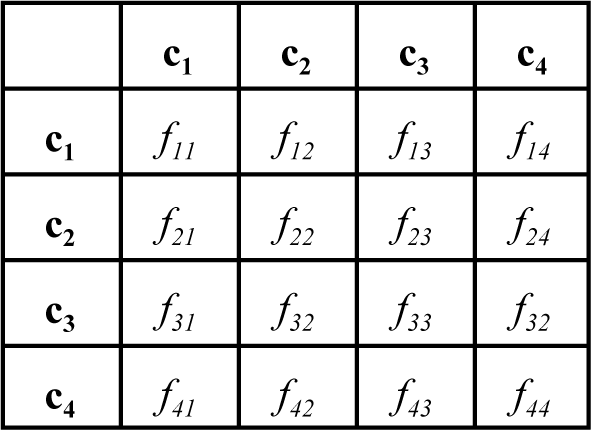
example of a raw edge length or frequency matrix **T_E_** and *C* = 4

In the raw **T_E_**, each, *f*_*ij*_ is the number of adjacent pixel pairs with the same (*i = j)* or different (*i* ≠ *j)* colors. If *i = j* then the transition was within a given color class. If *i* ≠ *j* then the transition was across the edge between two color classes. If each *f*_*ij*_ is divided by the sum of all of *f* then *f*_*ij*_ is the relative frequency of transition *i-j.* The diagonals (*f*_*ii*_) estimate the total area of each color, and the off-diagonals (*i* ≠ *j*) estimate the relative frequency or total length of the edges between colors *c*_*i*_ and *c*_*j*_. The upper and lower off-diagonals are just records of transitions in different directions, *i → j* and *j → i.* For subsequent analysis, the upper and lower off-diagonals should be combined as *f*_*k*_ = *f*_*ij*_ + *f*_*ij*_, where *k* = 1…E. and *E = C*(*C*-1)/2. In this example *E* = 6. For subsequent analysis use the *f*_*k*_ and ignore the diagonals, yielding the final version of **T_E_**, as in Table S2.

**Table S2.** example of a the final version of the edge transition matrix **T_E_**

| | $c1$ | $c2$ | $c3$ | $c4$ |
| --- | --- | --- | --- | --- |
| $c1$ | | | | |
| $c2$ | $f_a = f_{12} + f_{21}$ | | | |
| $c3$ | $f_b = f_{13} + f_{31}$ | $f_c = f_{23} + f_{32}$ | | |
| $c4$ | $f_d = f_{14} + f_{41}$ | $f_e = f_{24} + f_{42}$ | $f_f = f_{34} + f_{43}$ | |

In table S2 *f*_*k*_ where *k* = a,b,…f instead of 1,2̤6 to avoid confusion with *i* and *j*.

If **T_E_** was generated directly from a map of the edges, then the data will appear as in table S2 with only cells *f*_*k*_.

For either **T_E_** calculation, the *f*_*k*_ can be converted to frequencies by dividing by their total, **T**=∑ *f*_*k*_.

For **T_E_** generated either from transitions or directly from an edge map, there are potentially *E = C*(*C*-1)/2 edge frequencies or lengths *f*_*k*_. However, the larger the C, the more likely it is that some colors may not contact others, and, as a consequence, some of the *f*_*k*_ will be zero.

### 1.2 The AS Matrices, **T_SC_** and **T_SL_**

These are accumulated directly from calculating the Receptor Noise JND or signal/noise ratio ΔS for all possible combinations of the *C* colors. The matrix will resemble Table S, with *s*_*ij*_ = ΔS for colors *i* and *j* instead of *f*_*ij*_, but the diagonals will be zero and the upper and lower off-diagonals will be identical for a given *i* and *j*. In this case just take the lower off-diagonals (call them *s*_*k*_), giving a matrix in the same form as Table S2.

### 1.3 Analysing **T_E_, T_SC_** and **T_SL_**

For all three transition matrices, **T_E_, T_SC_** and **T_SL_**, rearrange their *f*_*k*_ and *s*_*k*_ into column vectors. This can be done easily with the reshape function in MATLAB or the as.vector function in R. For further analysis it is convenient to place the three vectors into a E × 3 data matrix with one row per transition class, as in table S3. See main text for how the data matrix is used.

**Table S3.** transition matrices from Table S2 converted into vectors

| $\mathbf{T}_{\text{E}}$ | $\mathbf{T}_{\text{SC}}$ | $\mathbf{T}_{\text{SL}}$ |
| --- | --- | --- |
| $f_a$ | $s_a$ | $s_a$ |
| $f_b$ | $s_b$ | $s_b$ |
| $f_c$ | $s_c$ | $s_c$ |
| $f_d$ | $s_d$ | $s_d$ |
| $f_e$ | $s_e$ | $s_e$ |
| $f_f$ | $s_f$ | $s_f$ |

## 2. MATLAB function to calculate weighted mean and weighted standard deviation

~~~
function [mn,sd]=WeightedMnSD(x,w)
% [mn,sd]=WeightedMnSD(x,w);
% Calculates the mean and SD for x data with weights w INPUT: x values
%        w       weights for     each    value
% Both x and w must be the same length
% Will automatically remove any rows with NaN in x
% Formulae from the DATAPLOT manual pages 2-66 to 2-67 at
% http://www.itl.nist.gov/div898/software/dataplot/refman2/ch2/weightsd.pdft=isnan(x);
if sum(t)>0 %remove NaN rows xx=x(t==0); ww=w(t==0); x=xx; w=ww; w=w/sum(w);
end;
n=length(x); n2=length(w); if n~=n2
mn=NaN; sd=NaN;
fprintf(1,'X and weights do not have same n\n'); return;
end;
sw=sum(w);      %sum of weights
swx=sum(x.*w); %sum of weights times x mn=swx/sw;       %weighted mean
nnz=sum(w(w>0)>0); %number of nonzero weights if nnz>1
s=0;
for i=1:n
s=s+w(i)*(x(i)-mn)^^l^2;
end;
sd=sqrt(nnz*s/((nnz-1)*sw)); %math.stackexchange.com else
sd=0;
end;
~~~

## 3. Gouldian Finch example in more detail

The analysis was done for two views of each morph, digitized from photographs. The two views may represent two different views as seen by conspecifics but in any case demonstrate the effects of different views of the same bird. The illustrations below are maps of the patch boundaries.

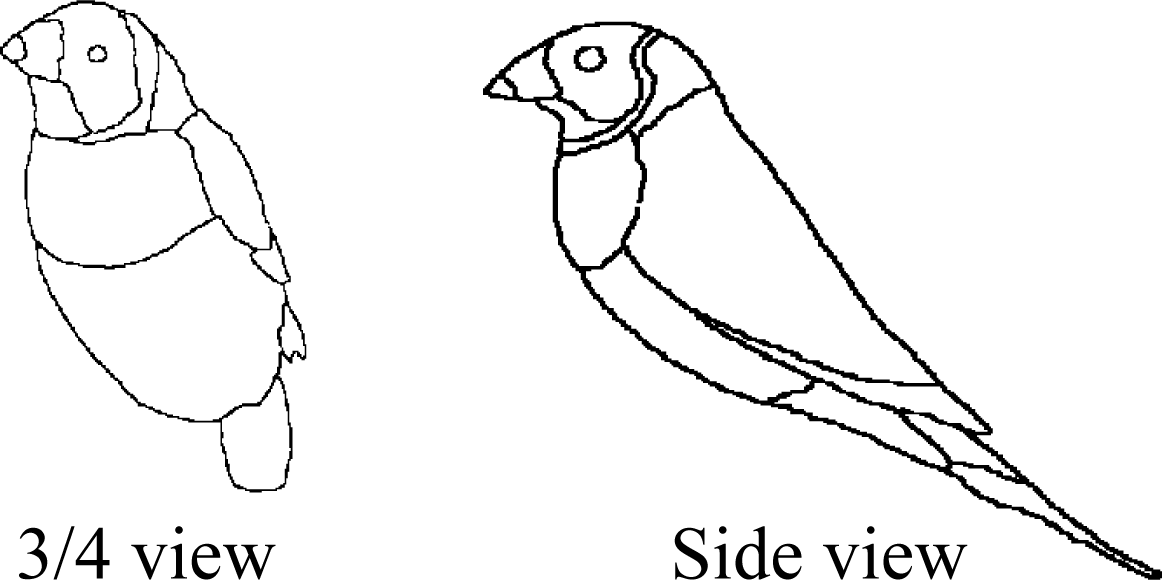

The following 4 pages show various combinations of morph (Black, Golden and Red), sex, and view.

Note that, for clarity, the boundary height (intensity) between the bird and the background is shown as zero. When seen against real background there would be fluctuations around the bird’s perimeter both above and blow the bird’s own patch boundary intensity.

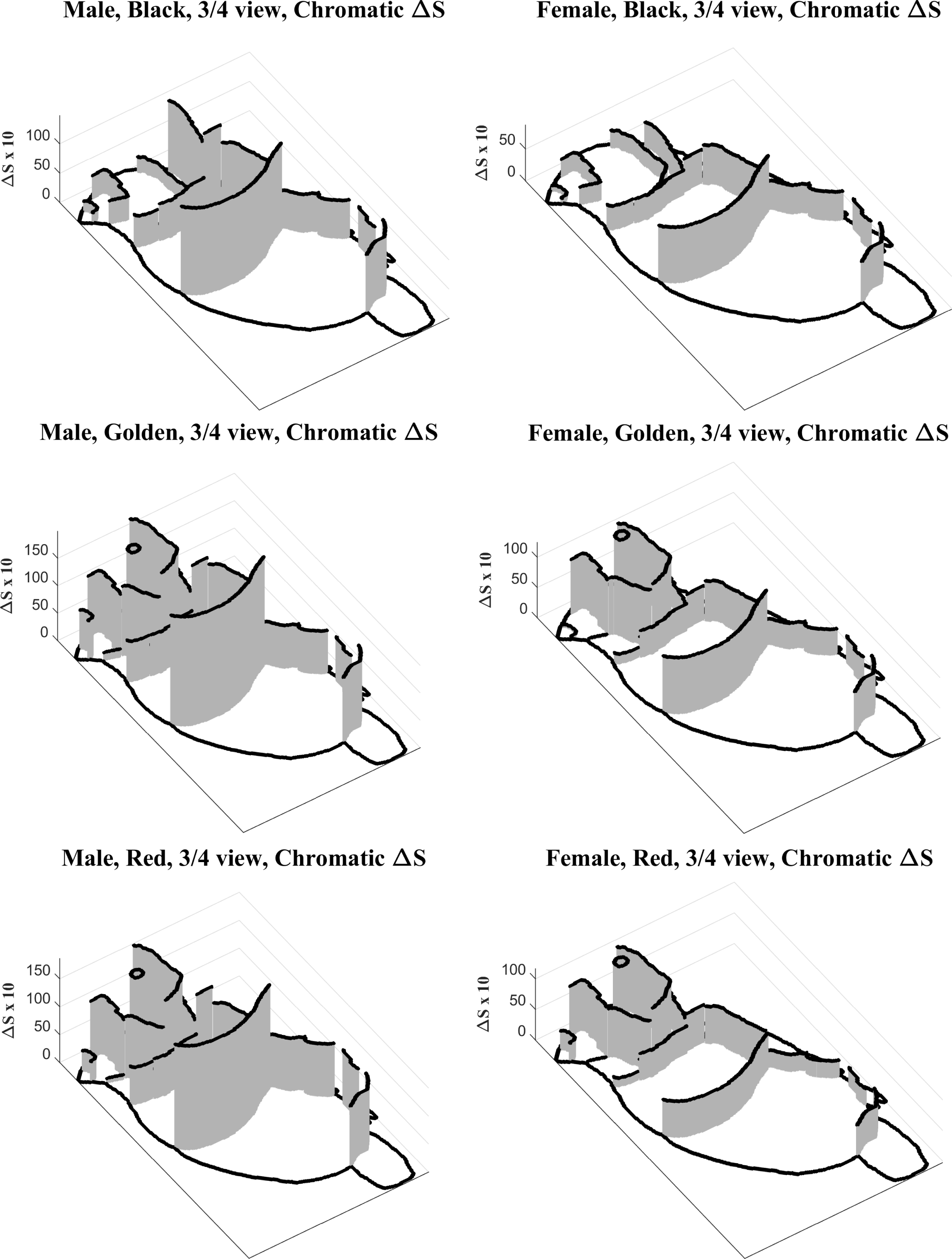

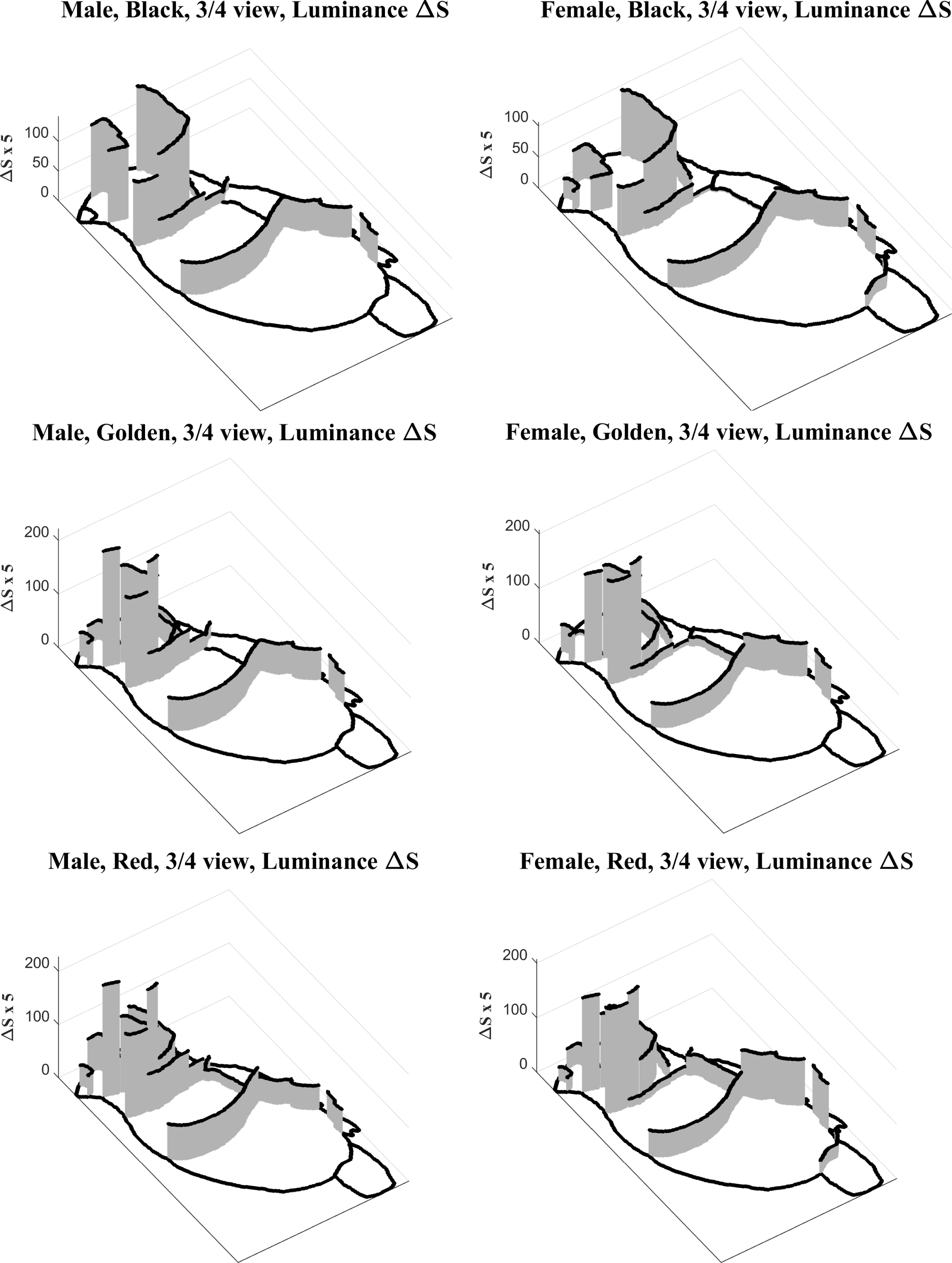

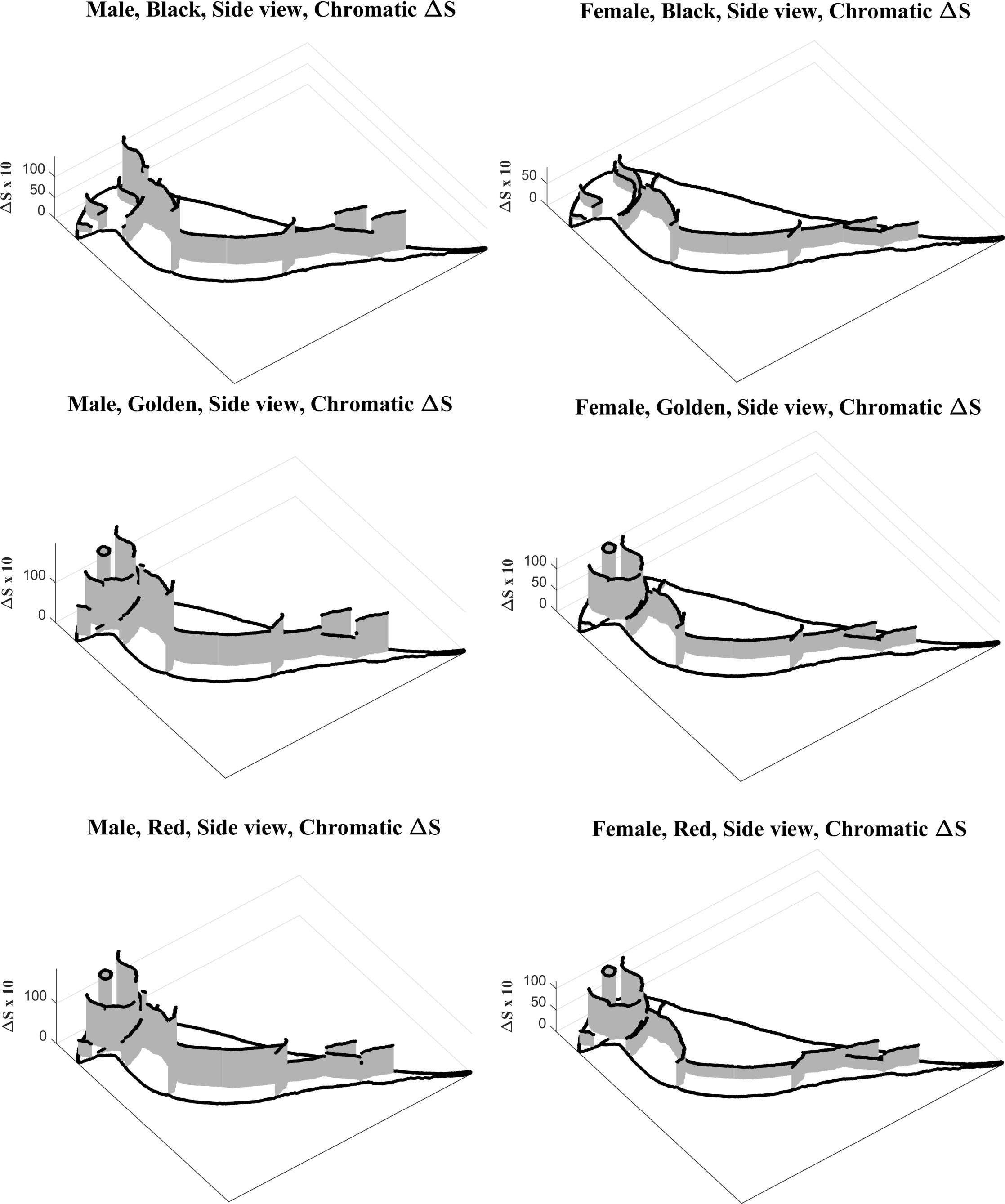

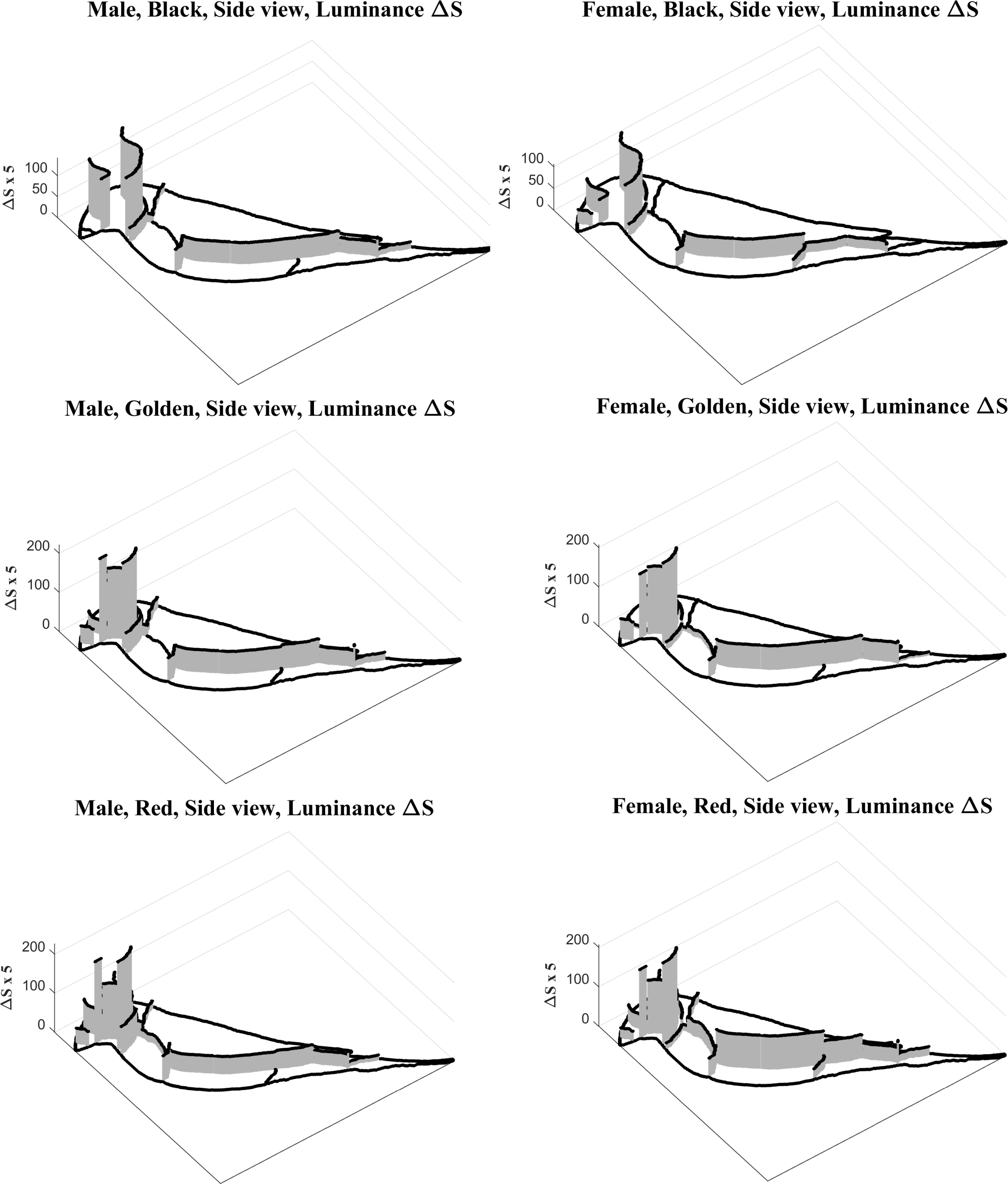

## Gouldian Finch statistics

### Correlation between chr and lum edge deltaS, point per pixels Excluding edges between body and background and eye

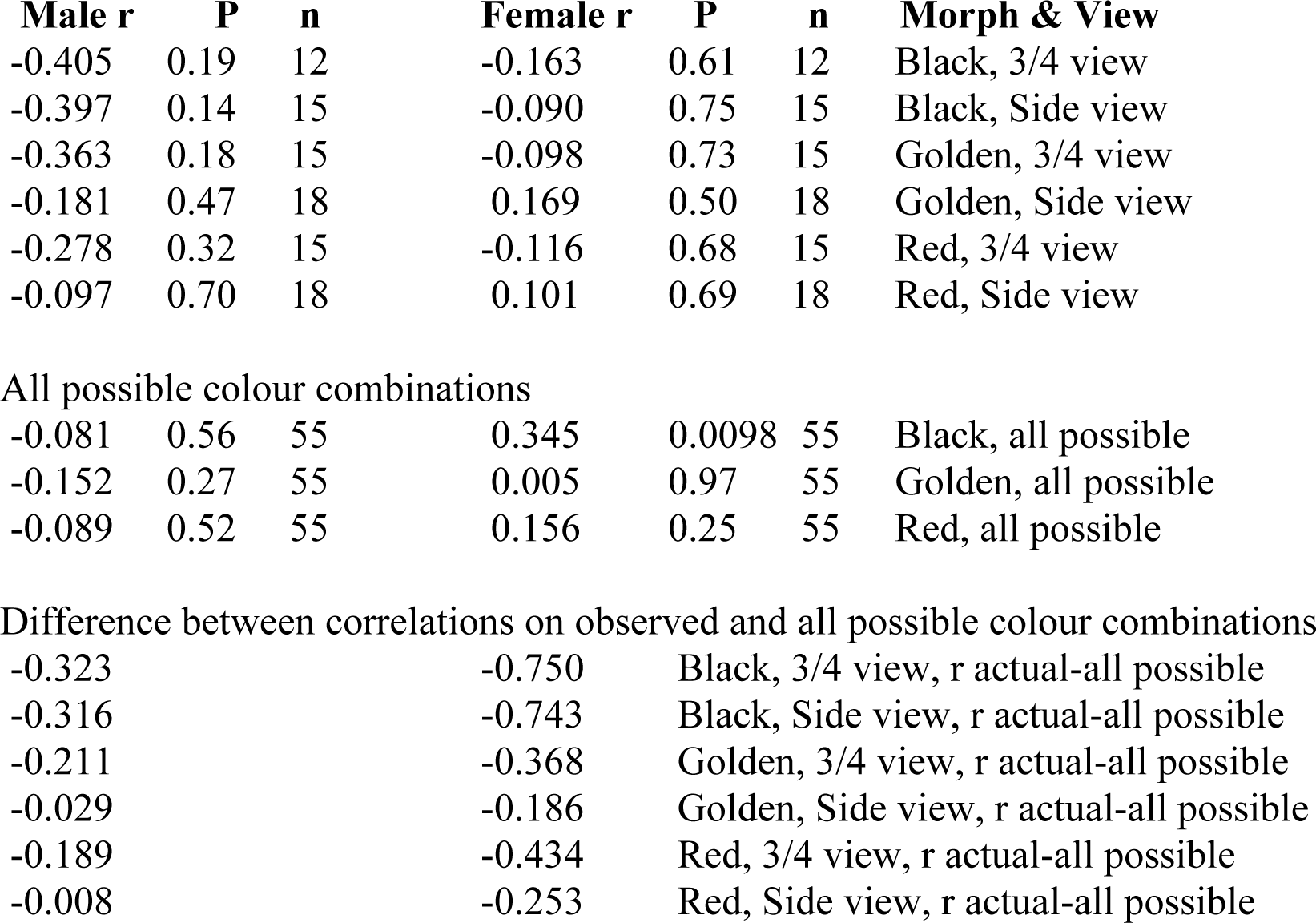

| Male r | P | n | Female r | P | n | Morph & View |
| --- | --- | --- | --- | --- | --- | --- |
| -0.405 | 0.19 | 12 | -0.163 | 0.61 | 12 | Black, 3/4 view |
| -0.397 | 0.14 | 15 | -0.090 | 0.75 | 15 | Black, Side view |
| -0.363 | 0.18 | 15 | -0.098 | 0.73 | 15 | Golden, 3/4 view |
| -0.181 | 0.47 | 18 | 0.169 | 0.50 | 18 | Golden, Side view |
| -0.278 | 0.32 | 15 | -0.116 | 0.68 | 15 | Red, 3/4 view |
| -0.097 | 0.70 | 18 | 0.101 | 0.69 | 18 | Red, Side view |
| All possible colour combinations |  |  |  |  |  |  |
| -0.081 | 0.56 | 55 | 0.345 | 0.0098 | 55 | Black, all possible |
| -0.152 | 0.27 | 55 | 0.005 | 0.97 | 55 | Golden, all possible |
| -0.089 | 0.52 | 55 | 0.156 | 0.25 | 55 | Red, all possible |

Difference between correlations on observed and all possible colour combinations
|  |  |  |
| --- | --- | --- |
| -0.323 | -0.750 | Black, 3/4 view, r actual-all possible |
| -0.316 | -0.743 | Black, Side view, r actual-all possible |
| -0.211 | -0.368 | Golden, 3/4 view, r actual-all possible |
| -0.029 | -0.186 | Golden, Side view, r actual-all possible |
| -0.189 | -0.434 | Red, 3/4 view, r actual-all possible |
| -0.008 | -0.253 | Red, Side view, r actual-all possible |

### Mean (*m*_*ΔS*_) and SD (*s*_*ΔS*_) of patch edge chromatic (cr) and luminance (lm) *ΔS*, weighted by edge lengths

Mean ( $m_{\Delta S}$ ) and SD ( $s_{\Delta S}$ ) of patch edge chromatic (cr) and luminance (lm) $\Delta S$ , weighted by edge lengths
| Cr $m_{\Delta S}$ | Cr $s_{\Delta S}$ | Lm $m_{\Delta S}$ | SD_Lm | Morph-gender-view |
| --- | --- | --- | --- | --- |
| 7.56 | 4.97 | 11.07 | 10.43 | Black, Male, 3/4 view |
| 5.71 | 4.25 | 7.84 | 9.29 | Black, Male, Side view |
| 4.49 | 2.64 | 8.55 | 6.53 | Black, Female, 3/4 view |
| 3.19 | 2.21 | 5.78 | 6.50 | Black, Female, Side view |
| 12.30 | 5.46 | 11.84 | 11.08 | Golden, Male, 3/4 view |
| 8.58 | 5.55 | 9.75 | 10.91 | Golden, Male, Side view |
| 6.70 | 3.43 | 9.90 | 10.41 | Golden, Female, 3/4 view |
| 4.77 | 3.57 | 8.33 | 9.91 | Golden, Female, Side view |
| 11.44 | 4.94 | 12.95 | 9.56 | Red, Male, 3/4 view |
| 7.96 | 4.95 | 9.68 | 10.10 | Red, Male, Side view |
| 5.75 | 2.98 | 11.80 | 9.30 | Red, Female, 3/4 view |
| 4.40 | 3.25 | 9.76 | 9.24 | Red, Female, Side view |

